# Towards the extended barcode concept: Generating DNA reference data through genome skimming of danish plants

**DOI:** 10.1101/2021.08.11.456029

**Authors:** Physilia Y. S. Chua, Frederik Leerhøi, Emilia M. R. Langkjær, Ashot Margaryan, Christina L. Noer, Stine R. Richter, Marlene E. Restrup, Hans Henrik Bruun, Ida Hartvig, Eric Coissac, Sanne Boessenkool, Inger G. Alsos, Kristine Bohmann

**Affiliations:** Section for Evolutionary Genomics, GLOBE Institute, Faculty of Health and Medical Sciences, University of Copenhagen, 1353 Copenhagen K, Denmark; Center for Evolutionary Hologenomics, University of Copenhagen, 1353 Copenhagen K, Denmark; Department of Biology, Faculty of Science, University of Copenhagen, 2200 Copenhagen K, Denmark; Department of Geosciences and Natural Resource Management, University of Copenhagen, 1958 Frederiksberg C, Denmark and Smithsonian Environmental Research Center, Smithsonian Institute, Edgewater 21037, Maryland, USA; LECA, Univ. Grenoble Alpes, Univ. Savoie Mont Blanc, CNRS, F-38000 Grenoble, France; Centre for Ecological and Evolutionary Synthesis (CEES), Department of Biosciences, University of Oslo, Blindernveien 31, NO-0371, Oslo, Norway; The Arctic University Museum of Norway, UiT – The Arctic University of Norway, N-9037 Tromsø, Norway

**Keywords:** Chloroplast, genome skimming, molecular studies, nuclear ribosomal DNA, plant DNA barcode, plastid genome

## Abstract

**Background:** Recently, there has been a push towards the extended barcode concept of utilising chloroplast genomes (cpGenome) and nuclear ribosomal DNA (nrDNA) sequences for molecular identification of plants instead of the standard barcode regions. These extended barcodes has a wide range of applications, including biodiversity monitoring and assessment, primer design, and evolutionary studies. However, these extended barcodes are not well represented in global reference databases. To fill this gap, we generated cpGenomes and nrDNA reference data from genome skims of 184 plant species collected in Denmark. We further explored the application of our generated reference data for molecular identifications of plants in an environmental DNA metagenomics study.

**Results:** We assembled partial cpGenomes for 82.1% of sequenced species and full or partial nrDNA sequences for 83.7% of species. We added all assemblies to GenBank, of which chloroplast reference data from 101 species and nuclear reference data from 6 species were not previously represented. On average, we recovered 45 genes per species. The rate of recovery of standard barcodes was higher for nuclear barcodes (>89%) than chloroplast barcodes (< 60%). Extracted DNA yield did not affect assembly outcome, whereas high GC content did so negatively. For the *in silico* simulation of metagenomic reads, taxonomic assignments using the reference data generated had better species resolution (94.9%) as compared to GenBank (18.1%) without any identification errors.

**Conclusions:** Genome skimming generates reference data of both standard barcodes and other loci, contributing to the global DNA reference database for plants.

## BACKGROUND

Accurate identification of plant taxa is integral to many molecular studies and has been used in studies including the identification of plants from pollen for hay fever forecasting [1], for detecting adulteration in herbal medicine [2], and authenticating wood samples for commercially traded timber trees [3]. Plants can also be molecularly identified from ancient sedimentary DNA (sedaDNA) samples to reconstruct past vegetation [4,5], environmental DNA (eDNA) samples such as water for water quality assessments [6], faeces for diet authentication of grazing livestock [7] or to reveal new ecological information about animals’ diet [8], and soil for biodiversity monitoring [9,10]. Accurate identification is essential for biodiversity monitoring studies as conservation efforts are dependent on obtaining species distribution information. Such identifications are made by comparing sequenced DNA extracted from samples to DNA reference databases [11,12]. As such, the DNA reference database is the foundation of molecular studies. Plant DNA reference data can be created through the sequencing of samples from taxonomically identified voucher specimens deposited in curated museum collections. The use of taxonomically verified specimens to generate DNA reference data reduces the risks of misidentification between taxa and DNA references [13–15].

Plant DNA reference databases traditionally consist of reference sequences, so-called barcode sequences, that have been generated through PCR amplification with universal primers designed to target specific regions of the chloroplast genes. These barcodes are short, conserved sequences of DNA that act as molecular identifiers for each species [12]. For plant reference data generation, the two most commonly used chloroplast barcodes are the ribulose-bisphosphate carboxylase large-chain (*rbcL*, ~654 bp) and maturase K (*matK*, ~889 bp) genes [16–20]. Even though these two genes have been proposed as standard barcodes for the identification of plant taxa [16], they have their limitations [21,22]. For example, *matK* does not amplify well in non-angiosperms, and the power of species discrimination between closely related species is generally insufficient for both barcodes [23]. For applications analysing plant taxonomic composition from highly degraded DNA or mixed templates such as eDNA samples, a chloroplast minibarcode *trnL* P6 loop intron (~10 to 143 bp) and a smaller nuclear barcode of the internal transcribed spacers (ITS), ITS2 (~221 to 260 bp), have been proposed as they are short enough to be amplified from degraded samples [11,17,24–27]. However, these barcodes also have limitations such as co-amplification of fungal DNA for ITS2 and typically poor species resolution for *trnL* P6 loop intron (33-93%% to species level [28,29]). This lack of consensus on which barcodes to use for plant identification has led to the suggestion of routinely using two or more barcodes for molecular plant identification [12,16,30–32].

The taxonomic resolution that can be obtained is directly related to the DNA reference databases; the more complete the reference database is, in terms of the number of species represented and the number of barcode regions available per species, the higher is the taxonomic resolution [33]. The need to design suitable primers for targeted barcode amplification and risks of PCR failures where the targeted barcodes are not amplified has led researchers to move away from generating single-barcode plant reference sequences and instead, generate the so-called ‘genome skims’ [34]. Genome skims, also known as ‘extended barcodes’, are generated using the genome skimming approach, which is low-coverage high-throughput shotgun sequencing of the total DNA extracted from a given plant taxa. This universal extended barcode for plants has been proposed by several researchers [34–37]. Moreover, as genome skimming circumvents the barcode PCR amplification step used to amplify and sequence single-barcodes, it allows for easier generation of DNA reference data from degraded samples with fragmented DNA such as old herbarium voucher specimens. Additionally, large amounts of genomic data are generated [13,14,34,38–40], guaranteeing resolution to below species level for most taxa [41]. This data can be used for the assembly of chloroplast genomes (cpGenomes) and nuclear ribosomal DNA (nrDNA) sequences within which plant barcodes and other gene loci can be recovered [13,34,42]. The ability of genome skimming to go beyond generating standard plant barcodes is highly advantageous as the resulting genomic data can be used for other applications including species delimitation[16,22,43], population genomics [44], development of capture probes [45–47], evolutionary and phylogenetic studies [21,34,40,41,48–51], and metagenomics eDNA studies [52–54]. Hence, genome skimming is increasingly seen as the way forward to build large-scale curated DNA reference databases from taxonomically identified voucher specimens [13,14].

DNA reference data from about 69% of described land plant species are underrepresented in global DNA reference databases in terms of number of species and barcode regions represented [55,56]. In GenBank, only an estimated 0.36% of the 350,000 plant species have complete cpGenomes reference data represented (Accessed 13 June 2021), and the estimate is even lower for nrDNA sequences [57]. To fill the gap in reference data represented in global reference databases, national DNA reference database projects utilising genome skimming to assemble full or partial cpGenomes have been created. These projects cover the regional floras of Australia (PILBseq project) [14], China [40], the European Alps and Carpathians (PhyloAlps) and Norway/polar regions (PhyloNorway) [13]. In the Australian PILBseq project, Nevill et al. retrieved cpGenomes from 96.1% of herbarium samples, and nrDNA sequences from 93.3% of samples [14]. Further, they retrieved the two standard plant barcodes, *rbcL* and *matK*, for ca. 93.3% of the samples. However, they encountered assembly issues that were specific to taxa with low GC content. Similarly in the PhyloNorway project, Alsos et al. showed assembly rates for full cpGenomes (67%) and full nrDNA sequences (86%), with an average coverage of 278x and 603x, respectively [13]. They showed that a sequencing depth of 90x or higher is required for assembly. Additionally, all three common plant barcodes (ITS2, *rbcL*, and *matK*) were retrieved for more than 90% of all genera, independent of material type (silica-dried fresh plant materials or herbarium materials). These projects show that genome skimming is a feasible method to generate reference data and that the assembly rate is high using both silica-dried fresh plant materials or herbarium materials.

In Denmark, of the around 2,800 extant vascular plant species recorded [58], most species do not have cpGenomes or nrDNA sequences available in GenBank. With the increasing number of molecular biodiversity monitoring studies of plants carried out in Denmark [59], there is a demand for access to a well-curated DNA reference database comprising reference sequences representative of the local flora. The availability of a curated national DNA reference database of Denmark’s flora is essential for establishing future DNA-based monitoring programs. For example, for studying Denmark’s largest orchid *Orchis purpurea*, which is rare and threatened [60], or for monitoring rare habitat-types like the rich fens. Further, other applications utilising genomic reference data can similarly benefit. To meet these needs, we generated genome skims of 184 Danish vascular plant species. These genome skims were used to assemble cpGenomes and nrDNA sequences for the Danish national DNA reference database (DNAmark). The 184 Danish plant species selected for genome skimming cover 45% of all Danish vascular plant families. As a guide to future genome-skimming projects, we evaluate, i) DNA yield and GC content in relation to assembly outcomes, ii) recovery rates of five common plant barcodes (*rbcL*, *matK*, *trnL*, ITS1 & ITS2), and iii) sequencing depth exploration for assembly coverage and optimal recovery of common plant barcodes. In addition, we evaluated the application of our database as compared to GenBank for use in metagenomics eDNA studies through *in silico* simulation of plant metagenomics datasets. This work creates the foundation for adding more plant species to the growing DNAmark reference database and hence, establishing the means for successful molecular identification of plants for biodiversity monitoring studies in the future.

## METHODS

### Species selection

A national DNA reference database for Danish species was established in 2017 (DNAmark) to create genome skims of 1000 species of animals, plants, and fungi observed in Denmark [61]. The 184 plant species sequenced in this study were selected by the DNAmark committee consisting of over 40 Danish taxonomists, researchers, and conservation officers in Denmark. The sampled plant species were selected based on broad phylogenetic coverage, relevance to Danish nature conservation, importance to future research projects, and on availability. A total of 198 leaf samples were sampled from either fresh specimen collected between 2017 to 2018 (n = 177), or from vouchered herbarium specimens (n = 21) stored at Herbarium C of the National History Museum of Denmark. Freshly collected specimens were deposited into the herbarium after sampling. The identity of all sampled specimens was confirmed by plant taxonomists at the Natural History Museum or other botanical experts, to reduce the likelihood of inaccurate sequence identification. The sampled specimens spanned 30 taxonomic plant orders, 54 families, 124 genera, and 184 species (Table S1_Supplementary Material 1). Of the 198 specimens, 82 were collected from the Capital Region, 10 from Mid-Jutland, 71 from Southern Denmark, and 35 from Zealand (Fig 1). All leaf samples were either dried in silica beads (Sigma-Aldrich 1-3 mm particle size) or stored frozen at −20 °C before DNA extraction.

**Fig 1:**
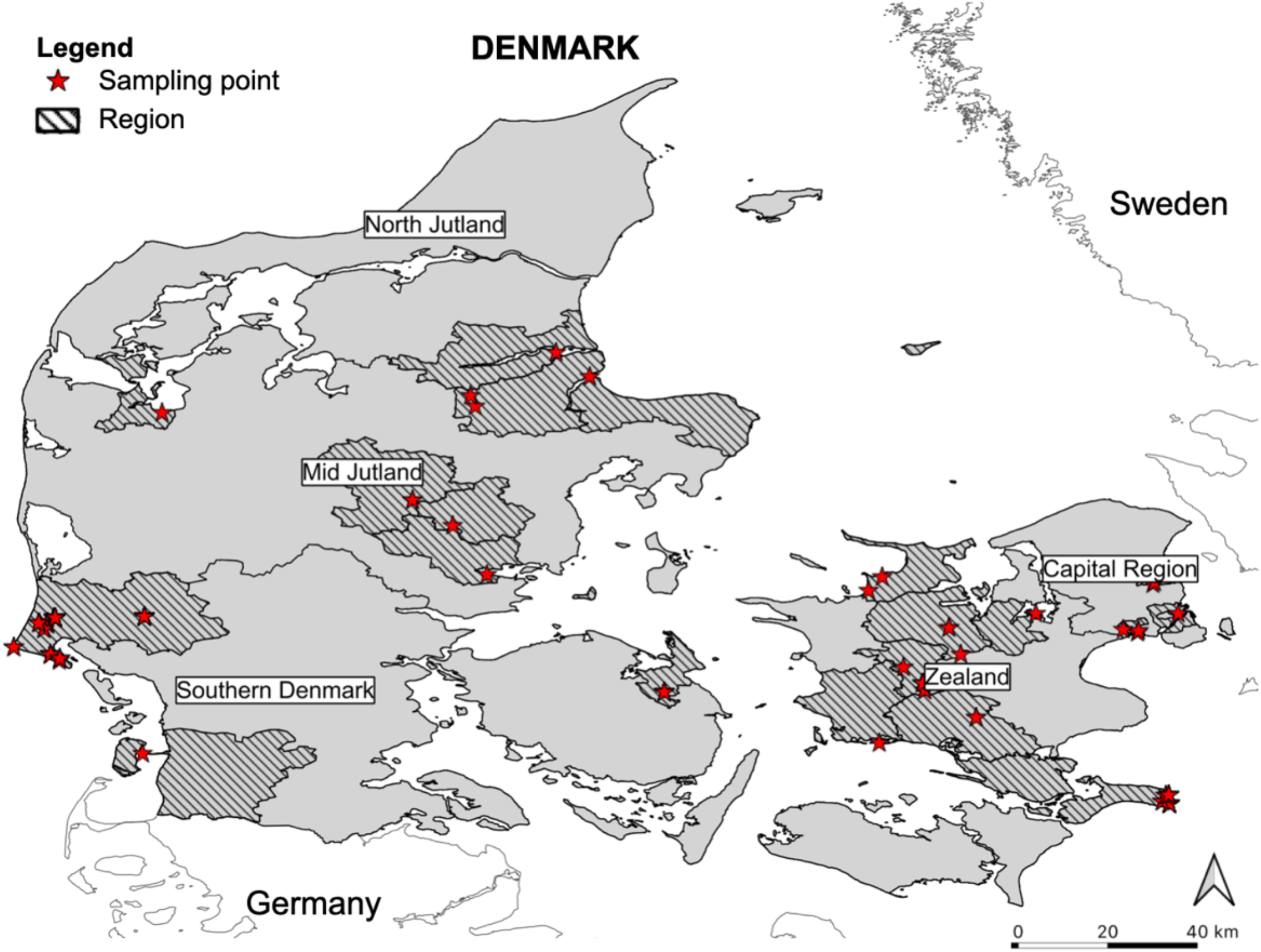
Location of all 198 vouchered herbarium specimens collected in Denmark from the Capital Region (n = 82), Mid-Jutland (n = 10), Southern Denmark (n = 71) and Zealand (n = 35) that were sequenced in this research.

### DNA extraction and sequencing

Samples were homogenized on a Qiagen Tissuelyser II with four to five 3 mm glass beads per tube for 40 seconds at 30 rpm (10 mg of silica-dried sample or 50 mg of frozen sample). DNA extractions were carried out on the KingFisher Duo Prime robot with Microtiter deep well 96 plates, using the Thermo Scientific™ KingFisher™ Pure DNA Plant Kit according to the manufacturer’s protocol (Thermo Scientific™ KingFisher™ Plant DNA Kit Instruction Manual Rev. 1.1), with the following modifications: Lysis buffer A was mixed with Polyvinylpyrrolidone (PVP) (0.09 g PVP to 4550 μL lysis buffer A) before adding to the samples. Prior to incubation, 25 μL of proteinase K was added to each sample and incubation time was increased to 1 hour. The deep 96-well plates were loaded according to the manufacturer’s instructions. To increase DNA concentration, elution volume was decreased to 80 μL. All DNA extracts were fragmented on a Covaris M220 Focused-ultrasonicator system aiming at an average of 475 (base pair) bp fragment size. Illumina sequence libraries were prepared using the Blunt End Multi Tube (BEMT) protocol [62,63] using 2 μL Illumina adaptors for each library. Before the index PCR, the libraries were purified using MAGBIO HighPrep™ PCR Clean-up System (1.6 times beads to library ratio) and eluted in 50 μL Qiagen EB buffer after 15 mins incubation at 37°C. The purified libraries were then screened on an MX3005 qPCR machine (Agilent) using 1 μL of 20x diluted purified libraries. The qPCR master mix contained 1x AmpliTaq Gold buffer (Applied Biosystems, USA), 2.5 mM MgCl2 (Applied Biosystems), 0.2 μM of each dNTPs (Invitrogen), 0.2 mM Bovine Serum Albumin (BSA) (Bio Labs), 0.1 U/μL TaqGold (Applied Biosystems), 0.8 μL SYBR Green I nucleic acid gel stain (S7563) (Invitrogen) with ROX Reference Dye (12223-012) (Invitrogen) (1:4 SYBR Green to ROX ratio with 2000 parts high-grade DMSO), and 0.2 μM of P5 and P7 indexed Illumina primers [64]. The qPCR was performed using the following conditions: 95°C for 10 mins, followed by 40 cycles of 95°C for 30 secs, 60°C for 1 min, 72°C for 1 min, and ending with a dissociation curve. The results of the qPCR were used to determine the optimal number of PCR cycles for each purified library for indexing PCR. The purified libraries were amplified using P5 and P7 indexed Illumina primers, using the same master mix as the qPCR, excluding SYBR Green and ROX Reference Dye. The indexed libraries were then pooled in equimolar concentration. Pooled indexed libraries had unique indices assigned to avoid false assignments during multiplexing, which occurs when leftover index primers randomly prime libraries, thus creating new indexed libraries [65,66]. PCR was performed using the following conditions: 95°C for 12 mins followed by 9, 15 or 23 cycles (determined by the qPCR for the specific libraries) of 95°C for 20 secs, 60°C for 30 secs, 72°C for 40 secs and ending with 72°C for 5 min. The pooled indexed libraries were sequenced using 150 bp PE (paired-end) chemistry on the Illumina HiSeq 4000 platform at the National High-throughput DNA Sequencing Centre (Copenhagen, Denmark) aiming at ca. 5 Gb per specimen.

### De novo assembly and annotation

Prior to assembly, adapter sequences were removed and low-quality bases (phred score=33) were trimmed from the 3’ end using *AdapterRemoval v2.3.1* [67]. Quality checks were carried out using *FastQC v0.11.5* after trimming [68]. For de novo assembly of cpGenomes and nrDNA sequences, we used the *ORGanelle ASseMbler* (*ORG.asm*) [69], a De Bruijn graph-based assembly program designed for this purpose. First, the paired reads were indexed with the *oa index* function, using a length cut-off which keeps 90% of the reads. During the indexing process, a minimum read length is determined and read pairs shorter than the minimum read length are discarded. Indices are also added to the reads for more efficient access to the reads in the subsequent assembly process. For the assembly, the *oa buildgraph* function was used with a seed from *Arabidopsis* to generate the assembly graph for both cpGenome and nrDNA. After the assembly graph was built, to extract the final assembled contigs, we used the *oa unfold* and *oa unfoldrdna* commands for the assembled cpGenome and nrDNA contigs respectively. Generation of circular contigs was first attempted for the cpGenome assembly. However, if that was not possible, one or more linear contigs were generated instead and used for downstream analysis [13]. We define assembly success as one or more contigs assembled, regardless if the assembly was full or partial, or if any of the standard barcode regions were assembled. When *ORG.asm* was unsuccessful in assembling the cpGenome, the de novo assembler *Novoplasty v.3.8.1* [70] was used instead with default parameters. For this, a reference sequence for assembly initiation (seed) was retrieved from the BOLD database [71] for each species. The seeds were chosen from the *rbcL or matK* barcode region. If the *rbcL or matK* barcode regions were not available for a particular species, *rbcL or matK* barcodes from a closely related species from the same genus were used instead. Assemblies of cpGenome and nrDNA sequences from both *ORG.asm* and *Novoplasty* were annotated using the *Organelle Annotator v.1.0.0* (*ORG.Annot*) [72].

### Quality control

The quality of the assembled sequences was checked by mapping the trimmed reads to the assembled sequence, using *Burrows-Wheeler Alignment v.0.7.17* (*BWA*), *bwa sampe* command [73]. *Samtools v.1.9* was used for sorting the mapped reads [74], and duplicates were removed using *Picard v.2.18.26* [75]. After removing duplicates, reads were realigned to the assembled sequence using *GenomeAnalysisTK v.4.1.2.0* (*GATK*) [76]. The resulting bam files were imported to *Geneious Prime v.2020.2* [77] and the assemblies were manually checked for errors. Following these quality control steps, files for GenBank submission were prepared using the *GB2sequin* online software [78].

### Sequencing depth exploration

To explore the sequencing depth impact on the cpGenome and nrDNA coverage as well as the proportion of common barcodes assembled, we attempted the assembly of cpGenome and nrDNA sequences using different sequencing depths. We only used a subset of our samples where all five barcode regions were assembled (*rbcL*, *matK*, *trnL*, ITS1 and ITS2) due to the computational-intensive nature of this exploration. As each sample represents one species, the subset of samples include nice species from nine families. These nine families (Amaranthaceae, Apiaceae, Caprifoliacea, Fabaceae, Orchidaceae, Orobanchaceae, Primulaceae, Rosaceae, and Salicaceae) were represented at least four times across all samples assembled for this study (Table S2_Supplementary Material 1). The adapter-removed and quality checked Illumina sequences from these subset were subsampled at an interval of 1 million PE reads (150 bp), until the maximum number of sequenced reads was reached. We used *seqtk v.1.3* [79] to carry out subsampling of reads, with the same random seed kept throughout. Assembly of these subsampled reads was carried out following the de novo assembly methods used for the DNAmark cpGenome and nrDNA sequences.

### *In silico* metagenomics application

To test the resolution that can be obtained from our DNAmark DNA reference database as compared to a global DNA reference database such as GenBank, we carried out *in silico* simulation of metagenomic reads using the software *InSilicoSeq* [80]. We simulated three Illumina metagenomic datasets consisting of plants based on the error model of a Hiseq instrument (125 bp PE, 1 million reads). Each dataset consists of cpGenome and nrDNA sequences of up to 23 plant species commonly found in Denmark (18 families and 22 genera) assembled in this study, where all five common barcodes were assembled (*matK*, *rbcL*, *trnL*, ITS1, ITS2) (Table S3_Supplementary Material 1). For each of the three simulated datasets, different numbers of species composition were used (dataset 1: 10 species, dataset 2: 15 species, and dataset 3: 23 species) (Table S4_Supplementary Material 1). Each of these three datasets was simulated twice to generate two replicates per dataset (dataset 1.1,1.2,2.1,2.2,3.1,3.2). Due to the simulation software used, for each replicate per dataset, the species composition would be identical but reads generated would be randomly selected from the cpGenome and nrDNA sequence input. From the study in Chua et al. [54], the authors highlighted that the use of whole-genome reference sequence for plant identification is currently not a feasible approach due to the lack of species representation. We have also calculated the species and genus representation of these five barcodes (96% species coverage, 100% genus coverage) as compared to utilising whole genomes from RefSeq (0% species coverage, 18% genus coverage) (Table S5_Supplementary Material 1). Therefore, for the *in silico* simulation, we have chosen instead to use the five common barcodes, which are more comprehensive and readily accessible.

We generated the GenBank DNA references for the five barcodes (downloaded March 2020, 65810 species present in all five databases) [55,83] according to Srivathsan et al. [52]. For the DNAmark DNA reference database used in the *in silico* testing, using a custom bioinformatics script we pulled out sequences corresponding to the five barcodes from the cpGenome and nrDNA sequences generated in this study, and used it as a separate DNA reference database for the matching of reads. For the taxonomic assignment of reads to each of these two databases (GenBank and DNAmark), we followed the steps outlined in Chua et al. [54]. The lowest common ancestor (LCA) algorithm was used for the assignment of taxonomic ranks [84]. Reads were identified to the lowest taxonomic rank based on the criteria of 98% sequence identity and 85 bp sequence overlap, using a python script *readsidentifier* (v.1.0) [52]. We only kept identification if both forward and reverse reads matched at a given taxonomic level.

Additionally, a second round of identification parameters utilising specific barcode combinations was used for accurate taxonomic identification at any given level. Species-level identifications were retained only when reads were mapped to any of the following barcode combinations: at least two or more chloroplast barcodes, both nuclear barcodes, or any chloroplast barcode(s) with any nuclear barcode(s). If species-level identification were not achieved, genus-level identifications were retained with any of the following barcode combinations: at least two or more chloroplast barcodes, one or both nuclear barcodes, or any chloroplast barcode(s) with any nuclear barcode(s). If neither species nor genus-level identification was possible, family level identifications were retained with any of the following barcode combinations: at least two or more chloroplast barcodes, one or both nuclear barcodes, or any chloroplast barcode(s) with any nuclear barcode(s). If neither species, genus, nor family level identification was possible with the specified barcode combinations, the remaining reads were discarded.

### Statistical Analyses

All statistics were carried out in *Rstudio v.3.6.2* [81]. To test for the effect of DNA yield and GC% content on assembly outcomes, we used the R base package Stats to conduct Shapiro tests of normality. As the Shapiro test of normality significantly deviated from normality for DNA yield (cpGenome: p=0.953 nrDNA: p=0.048) and GC% content (cpGenome: p=0.016, nrDNA: p=0.005), we carried out the tests of significance using the average values through the Wilcoxon rank-sum test. For the test of correlation between sequencing depth and assembly coverage, after testing for normality, we carried out a non-parametric Spearman’s Rho correlation test.

## RESULTS

A total of 184 plant species from 198 voucher specimens were sequenced (Table S1_Supplementary Material 1) (Fig 2). These covered 68% of orders, 45% of families, 20% of genera, and 6.6% of species of the approximately 2800 plant species recorded from Denmark. From these voucher specimens, a total of 161 partial chloroplast genomes (cpGenomes) (81.3% of specimens, 82.1% of species) and 165 nuclear ribosomal DNA (nrDNA) sequences (83.3% of specimens, 83.7% of species) were assembled (Table 1). For each voucher specimen, approximately 1.1% of the reads were mapped to cpGenomes (SD = 1.1%), and 0.1% were mapped to nrDNA sequences (SD = 0.1%). From our study, we added new reference data to GenBank of which cpGenomes from 101 plant species and nrDNA sequences from 6 plant species were previously not represented (Table S6 to S8_Supplementary Material 1). For most of the plant families, cpGenomes (94%) and nrDNA sequences (91%) were assembled with only Convolvulaceae (n=1) not having any contigs assembled (Figure S1_Supplementary Material 2).

**Fig 2:**
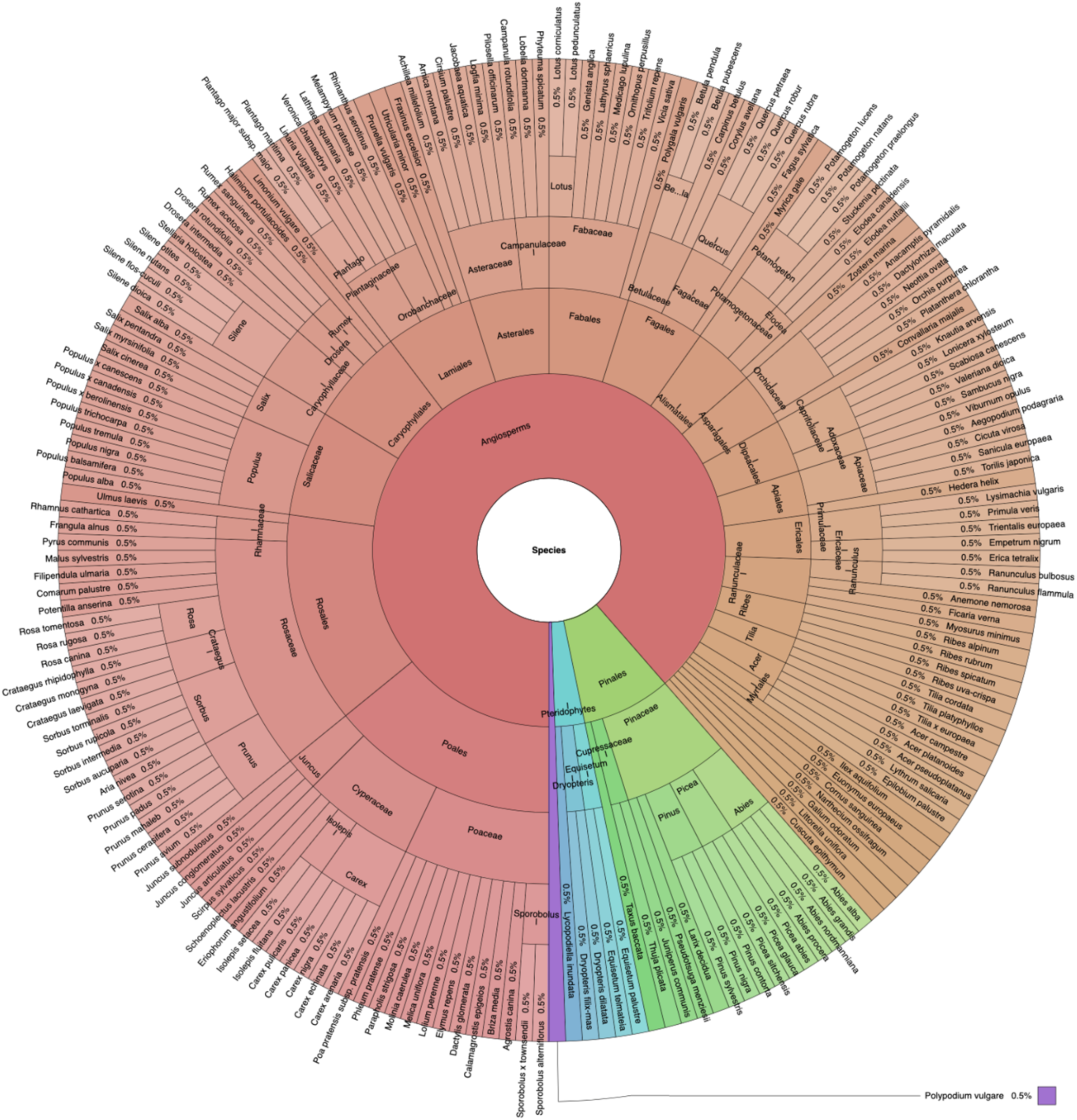
The taxonomic distribution of the DNA reference data consisting of plastid and nuclear ribosomal DNA reference data generated from genome skimming of 184 Danish plant species.

**Table 1:** Details of chloroplast genomes (cpGenomes) and nuclear ribosomal DNA (nrDNA) sequences assembled in this study

| Assembly details |  |
| --- | --- |
| Average PE reads before trimming per specimen: | 14,513,039bp |
| Average PE reads after trimming per specimen: | 14,505,561bp |
| Average sequence length per specimen: | 131bp |
| Number of partial cpGenomes assembled: | 161 partial cpGenomes |
| Average coverage of cpGenomes assembled: | 327x |
| Length range of cpGenomes assembled: | 1008 to 445384 bp |
| Average length of cpGenome assembled: | 48472.5bp |
| Number of nrDNA sequences assembled: | 165 nrDNA sequences |
| Average coverage of nrDNA sequences assembled: | 2595x |
| Length range of nrDNA sequences assembled: | 1829 to 10202 bp |
| Average length of nrDNA sequences assembled: | 5126bp |

Assembly outcomes of cpGenomes were not affected by the extracted DNA yield (Successful assemblies: average=48.4 ng/μL, SD = 60.3 ng/μL, unsuccessful assemblies: average=39.2 ng/μL, SD = 36.9 ng/μL) (Mann-Whitney: p=0.953). However, extracted DNA yield was higher for assembled nrDNA sequences (average 48.4 ng/μL, SD=58.7 ng/μL) than for unassembled ones (average = 32.4 ng/μL, SD=33.7 ng/μL) (Mann-Whitney: p=0.048). Additionally, differences in GC% content (average = 40, SD = 4.2) affected the assembly outcomes for both cpGenomes (Successful assemblies: average =39.5 ng/μL, SD=3.8 ng/μL, unsuccessful assemblies: average=41.7 ng/μL, SD=5.1 ng/μL) (Mann-Whitney: p=0.016) and nrDNA sequences (Successful assemblies: average =39.6 ng/μL, SD=3.7 ng/μL, unsuccessful assemblies: average=43 ng/μL, SD=5.7 ng/μL) (Mann-Whitney: p=0.005), with increasing GC% content negatively impacting assembly (Fig 3). We recovered on average 40 (SD = 35) genes for each assembled partial cpGenome, and 5 genes (SD = 0.9) for each assembled full or partial nrDNA sequences. The types of genes recovered consist of coding sequences (CDS), rRNA and tRNA (Table S9_Supplementary Material 1). The large subunit ribosomal ribonucleic acid (LSU rRNA) was the most recovered nrDNA gene, found in 78.3% of all sequenced samples. For cpGenome, the adenosine triphosphate synthase subunit beta (atpB) was the most recovered gene, found in 40.9% of all sequenced samples. Per barcode region, the rate of recovery for the five barcode regions were; *matK* at 49.7%, *rbcL* 58.4%, *trnL* 34.8%, ITS1 89.1%, and ITS2 89.7% (Figure S2_Supplementary Material 2). Chloroplast barcodes had a poorer rate of recovery (< 60%) than nuclear barcodes (> 89%).

**Fig 3:**
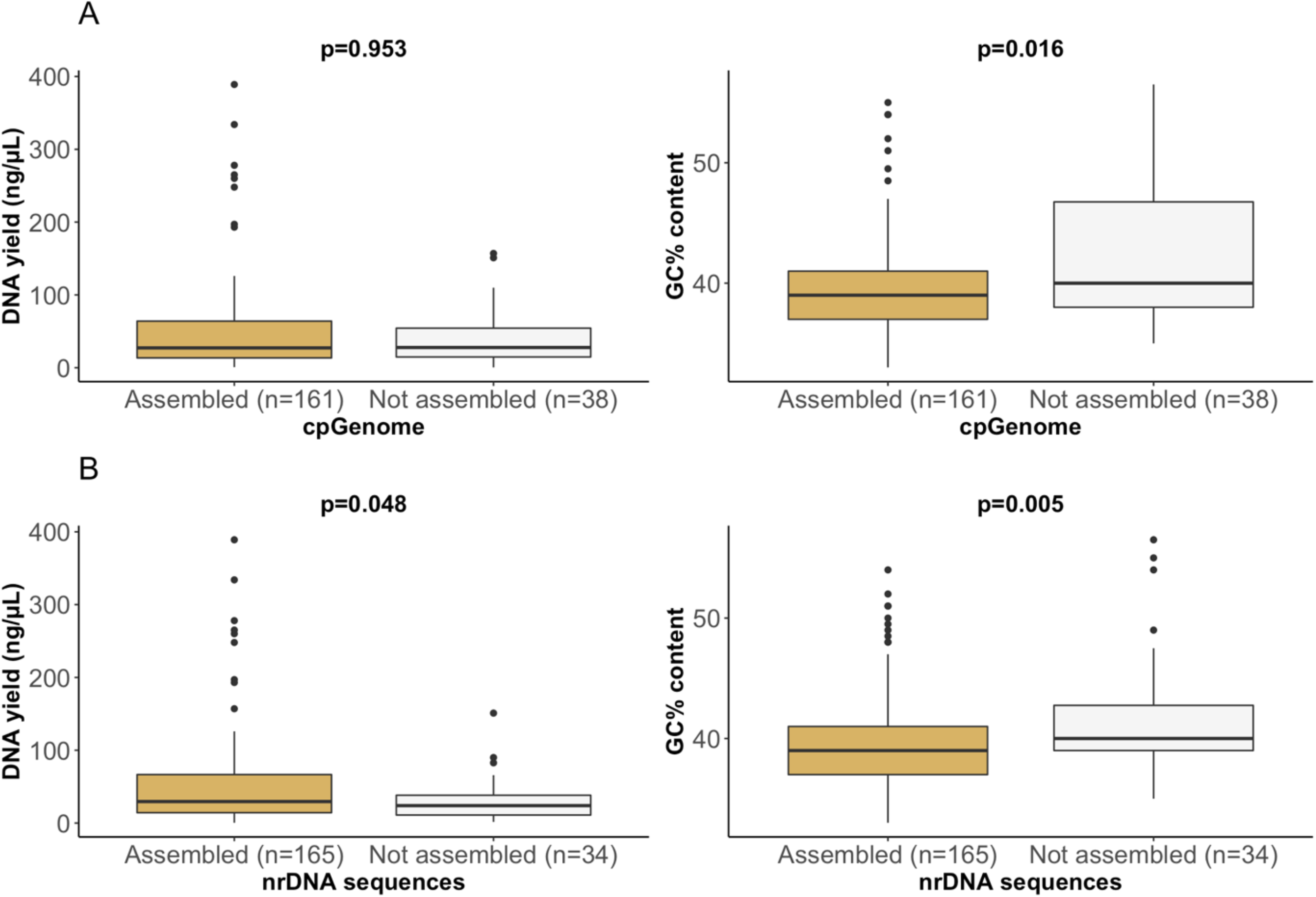
Box plot showing the distribution of DNA yield and GC% content in assembled (brown) and not assembled (grey) **a)** cpGenomes and **b)** nrDNA sequences. The p-values indicate the results of the Wilcoxon rank-sum test for differences in the average.

### Sequencing depth exploration

There was a weak positive correlation between sequencing depth and assembly coverage for cpGenome assembly (Spearman’s rho: 0.275, p=0.005), and a moderate positive correlation (Spearman’s rho: 0.567, p<0.001) for nrDNA sequences. For the assembly of barcodes at each sequencing depth, only 66.7% of sampled species had all three chloroplast barcodes (*matK*, *rbcL*, and *trnL*) assembled at 5 PE million reads (Fig 4). All three chloroplast barcodes were only assembled at the maximum sequencing depth (23 million bp) for *Melampyrum pratense*. Whereas for the nuclear barcodes (ITS1 and ITS2), we were able to assemble both barcodes in all nine sampled species using only 1 million PE reads, with a minimum coverage of 41x (Table S10_Supplementary Material 1).

**Fig 4:**
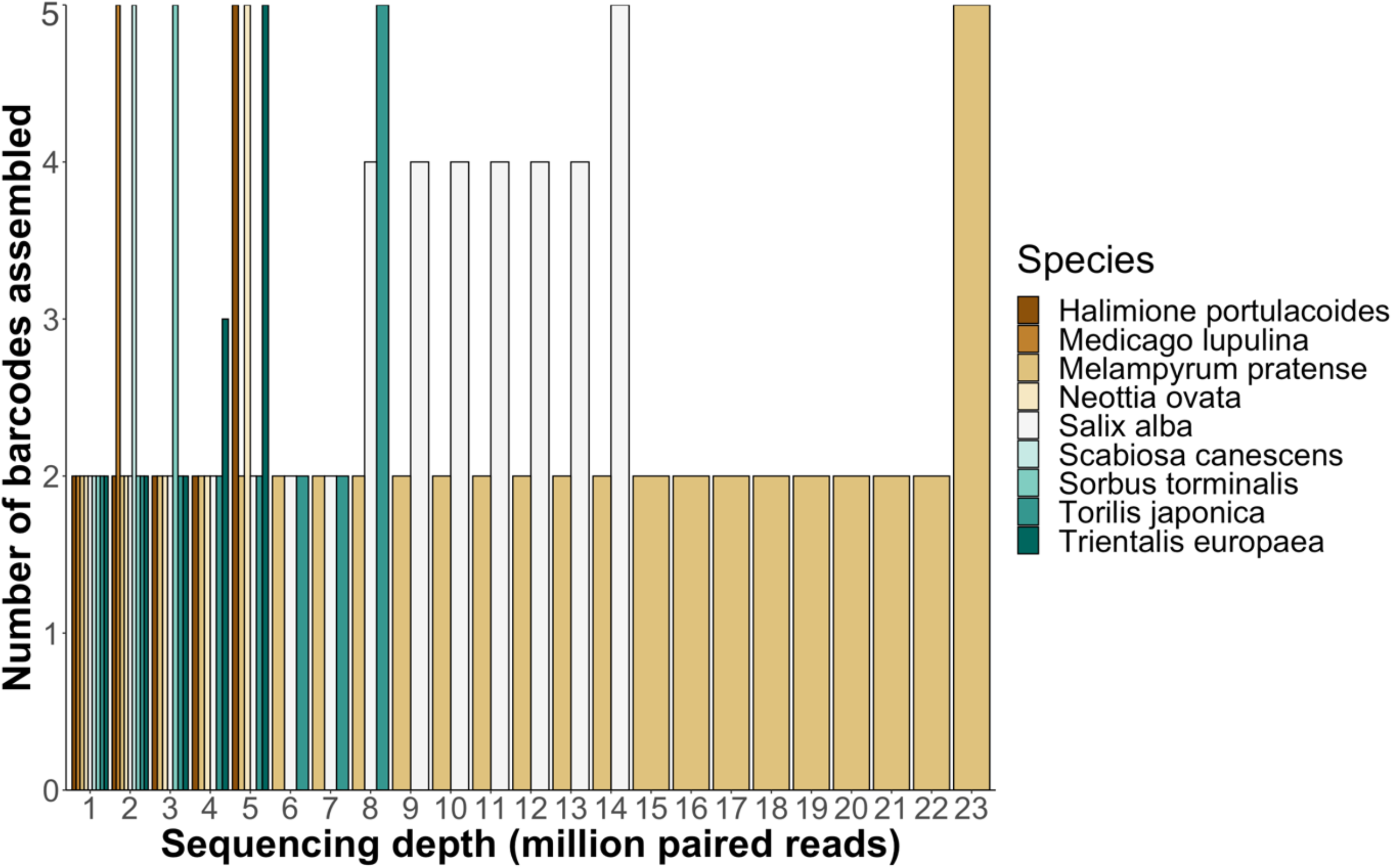
Number of assembled barcodes (*rbcL*, *matK*, *trnL*, ITS1 & ITS2) at each sequencing depth (PE) used in the sequencing depth exploration for nine species of plants. Subsampling at an interval of 1 million PE reads (150bp) for each species was done until the maximum number of sequenced reads were reached, or when all five barcodes were assembled.

### *In silico metagenomics* application

For the *in silico* metagenomics application, between 0.005% to 0.011% (average = 0.008%, SD = 0.003%) of metagenomic reads were mapped to plants for both databases (Table S11_Supplementary Material 1). After the second round of identification utilising specific barcode combinations, between 0.003% to 0.007% (average = 0.03%, SD = 0.03%) of reads were mapped to GenBank, while 0.005% to 0.01% (average = 0.007%, SD = 0.002%) of reads were mapped to our generated genome skimming reference data. The increase in the proportion of reads mapped to our generated genome skimming reference data as compared to GenBank was between 0.15% to 0.32% (average = 0.23%, SD = 0.06%) (Table 2). For GenBank, between 0.002% to 0.005% (average = 0.003%, SD = 0.001%) of reads were discarded while for DNAmark, <0.001% of reads were discarded. Of the mapped reads to plants, up to 47.3% of reads mapped to GenBank were not useful for identification (mapped to higher taxonomic levels, did not have any assigned taxonomy level, or did not fulfil barcode combination criteria). By contrast, of the reads mapped to our generated genome skimming reference data, a smaller proportion of reads (<12.5%) was not used for identification. Inaccurate identifications were observed only for matches to the GenBank database (between 5.6% to 15.6%, average = 8.3%, SD = 3.7%) and not to our generated genome skimming reference data (Table S12_Supplementary Material 1). When we increased the diversity of plant species used in the *in silico* dataset from 10 species to 23 species, the proportion of false positives utilising the GenBank database increased from 5.6% to 15.6%.

**Table 2:** Details of each simulated metagenomic dataset in the *in silico* tests when utilising reference data from either i) GenBank or ii) the generated reference data from this study

|  | Number of reads | Database used | Mapped reads (raw) | Proportion of mapped reads (raw) % | Assigned reads | Proportion of assigned reads % | Discarded reads | Proportion of discarded reads % | Proportion of discarded reads/mapped reads % | Proportion increase in mapped reads as compared to GenBank % |
| --- | --- | --- | --- | --- | --- | --- | --- | --- | --- | --- |
| Dataset 1.1 | 999990 | GenBank | 49 | 0.005 | 31 | 0.003 | 18 | 0.002 | 36.7 | NA |
|  |  | DNAmark | 50 | 0.005 | 46 | 0.005 | 4 | <0.001 | 8 | 0.15 |
| Dataset 1.2 | 999988 | GenBank | 50 | 0.005 | 31 | 0.003 | 19 | 0.002 | 38.0 | NA |
|  |  | DNAmark | 50 | 0.005 | 46 | 0.005 | 4 | <0.001 | 8 | 0.15 |
| Dataset 2.1 | 999974 | GenBank | 67 | 0.007 | 39 | 0.004 | 28 | 0.003 | 41.8 | NA |
|  |  | DNAmark | 72 | 0.007 | 63 | 0.006 | 9 | <0.001 | 12.5 | 0.24 |
| Dataset 2.2 | 999982 | GenBank | 74 | 0.007 | 39 | 0.004 | 35 | 0.004 | 47.3 | NA |
|  |  | DNAmark | 72 | 0.007 | 63 | 0.006 | 9 | <0.001 | 12.5 | 0.24 |
| Dataset 3.1 | 999954 | GenBank | 110 | 0.011 | 72 | 0.07 | 38 | 0.004 | 34.5 | NA |
|  |  | DNAmark | 111 | 0.011 | 98 | 0.01 | 13 | 0.001 | 11.7 | 0.26 |
| Dataset 3.2 | 999976 | GenBank | 114 | 0.011 | 68 | 0.07 | 46 | 0.005 | 40.4 | NA |
|  |  | DNAmark | 111 | 0.011 | 100 | 0.01 | 11 | 0.001 | 10 | 0.32 |

Per dataset, the taxa identified for both replicates were the same regardless of the database used. Matches to the GenBank database had between 10% to 26.7% species resolution (average = 18%, SD = 6.8%), 68.2% to 73.3% genus resolution (average = 70.5%, SD = 2.1%), and 86.7% to 90% family resolution (average = 88.5%, SD = 1.4%). In all datasets, most taxa were recovered minimally to the family level, except for Amaranthaceae and Sapindaceae even though the corresponding sequences were present in the database. For matches to our generated genome skimming data, species resolution was between 91.3% to 100% (average = 94.9%, SD = 3.7%), genus resolution was between 93.3% to 100% (average = 96.3%, SD = 2.6%), and at family level resolution was 100% (Table S13 and S14_Supplementary Material 1). Most taxa were retrieved except *Arnica montana* (both genus and species level) and *Sorbus torminalis* (species level).

## DISCUSSION

The reference data generated in this study included a broad coverage of Danish plant families and species important to Danish conservation and research. From genome skimming 184 species, we retrieved partial chloroplast genomes (cpGenomes) from 101 species and nuclear ribosomal DNA (nrDNA) sequences from 6 species that were previously not represented in GenBank. This outcome is important for two reasons. First, it corroborates the findings of three other national large scale genome skimming projects, showing that genome skimming is a suitable method to generate large amounts of genomic data from plants [13,14,40]. Second, it highlights the potential of genome skimming in adding new information of underrepresented taxa (i.e. Asteraceae, Orchidaceae) to existing databases [56]. With four regional flora databases now built using this technique, we have moved from a ‘proof of concept’ phase on the feasibility of utilising genome skimming as a viable approach for obtaining genome assemblies, to demonstrating that this approach can be readily replicated at a large scale, or even in smaller laboratories with limited manpower. Given that DNA reference data from most plant species (~69%) is still underrepresented in global DNA reference databases [55,56,82], the development of regional flora databases should be prioritised and genome skimming would be an effective method to generate such databases. Filling the reference data gap of underrepresented species in reference databases is particularly important to molecular-based biodiversity studies, where unidentified sequences due to missing information in the DNA reference databases results in inadequate species detection and monitoring [83]. Additionally, the genomic data generated in this study has many applications, including metagenomic analysis of environmental DNA (eDNA) samples and phylogenetic studies.

Based on sequencing costs, even though genome skimming was five times more expensive than sequencing the two standard barcodes alone, we assembled on average 45 genes for each sample. As argued in a genome skimming study for utilising genome skimming over barcode-based sequencing [13], the ability to simultaneously retrieve complete or partial cpGenome and nrDNA sequences pays off compared to sequencing individual barcodes. Even if difficult regions like *matK* are not assembled through genome skimming, other taxonomically informative barcodes or loci could be assembled [34], reducing the risk of failed assemblies when sequencing only single barcodes. More importantly, no single barcode region can be used to successfully resolve species-level identification across all plants, and genome skimming gives the added advantage of allowing more regions to be targeted. This will be more cost-effective in the long run, especially when resequencing is required for failed single barcode sequencing. Additionally, the majority of the work was carried out by only two technicians within a short duration of time, indicating that this approach could also be carried out by laboratories with limited manpower and time [13]. However, one drawback was that the bioinformatic pipelines used for assembly were computationally heavy, requiring on average five days run time per sample per genome type with at least 10 CPUs and 30Gb of memory. Thus, an assembly of this scale (>50 species) would not be computationally attainable without the use of a dedicated server, which could be a limiting factor for smaller laboratories with limited funds. This issue could be circumnavigated through collaborations with other laboratories capable of computing large amounts of data.

### Rate of assemblies and sequencing depth exploration

The relatively-high rate of assemblies (82.1% for cpGenome and 83.7% nrDNA) across a broad range of plant families and genera in our study serves as a guide for future genome-skimming projects. Differences in assembly rate between plant families could be an avenue for future exploratory work which we were unable to do due to few representatives collected per family in our study. However, there appears to be some differences that would require attention going forward such as the lower rate of assembly in the Gymnosperms which typically have huge genomes, and some of the water plants (Hydrocharitaceae and Potamogenaceae). Our recovery rates for the two standard plant barcodes *matK* (49.7%) and *rbcL* (58.4%) were similar to those obtained in other barcode-based sequencing studies [84]. However, this is lower than the other two genome skimming studies [13,14], where they had at least 93% recovery rate for chloroplast markers at lower coverage (30 to 150x, and 280x, respectively). This could be due to short fragments from our study that were around 250 bp on average used for assembly, which makes it a challenge to assemble reads. Improvements in the laboratory procedures to include longer fragments will be necessary for the success of future genome skimming studies, for better assembly success. Other than the two standard barcodes, we retrieved other taxonomically informative barcodes (*trnL*, ITS1, and ITS2) commonly used in the molecular identification of plants through genome skimming. We evaluated extracted DNA yield and GC content in relation to assembly outcomes. We showed that even though DNA yields were low following DNA extraction of voucher specimens, this did not affect the assembly outcomes of cpGenomes. This result is similar to a genome skimming study of Australian plant species, showing that DNA yield did not affect their assembly outcomes [14]. However, from our study, unassembled specimens were associated with samples with high GC content (Fig 3). This made assemblies especially challenging for species with high GC content that had fragmented DNA and for which shorter reads were therefore generated (<300 bp). Genomic regions with low GC contents can similarly be problematic [14], due to low coverage sequencing in these areas which leads to fragmented assemblies [85]. Future genome skimming of taxa with high/low GC content could increase sequencing depth to increase the amount of data for assembly [85], use reads from long-read sequencing [14], or utilise a combination of assemblers that could deal with assembling short reads.

Based on the sequencing depth exploration, the correlation between sequencing depth and assembly coverage was stronger for nrDNA sequences as compared to cpGenomes. This could be due to the smaller sizes of nuclear ribosomal regions, thus making it easier to assemble with lesser reads as compared to cpGenomes. For the recovery of barcodes, our results showed that despite being present in lower copy numbers than plastid genes [21], a low sequencing depth of 1 million PE reads was sufficient to retrieve both nuclear barcodes with minimal coverage of 41x. This can act as a guide for future studies, particularly if the aim is to only retrieve nrDNA sequences as small amounts of sequence data is sufficient to assemble high-quality nrDNA sequences and nuclear barcodes. At this sequencing depth, more samples can be sequenced to produce high-quality nrDNA sequence assemblies to fill the gap in the DNA reference databases used for plant identification. This is because nrDNA sequences are underrepresented in DNA reference databases as compared to chloroplast sequences [86]. In contrast, chloroplast barcodes required a higher sequencing depth to be assembled (66.7% assembly at 5 million PE reads on average). Even though this observation is only based on nine species, we can extrapolate this result for most plant species, due to the average larger sizes of chloroplast barcodes (*trnL-F* ~994 bp, *rbcL* ~654 bp, *matK* ~889 bp) over nuclear barcodes (ITS1 ~278 bp, ITS2 ~250 bp) [22,87]. Hence, for optimal use of genome skimming to generate chloroplast genome assemblies and barcodes, we recommend that sequencing depth should ideally aim at minimally 100x coverage.

### Utilisation of reference DNA generated

When we compared the utility of using a broader global DNA reference database (GenBank) with the reference DNA generated from our study for molecular plant identification in metagenomics analysis, species resolution with GenBank was typically poorer at less than 27% even though there is a 96% species coverage in the database (except for *Scabiosa canescens*). On average, there were also about 8.7% of identification errors. Additionally, not all barcodes were well represented in GenBank across all the species used in the *in silico* application. For example, the corresponding *trnL* barcode sequence is only represented for 65% of the species. Comparatively, utilising DNAmark yielded at least 91% species resolution with no identification errors. This increment of minimally five times higher species resolution with no misidentifications, despite the smaller number of species represented by our genome skimming reference data (184 species, GenBank 65810 species), shows that using a larger, broader, DNA reference database reduces species resolution [24,28,88]. In a broad global DNA reference database with many more taxa represented for a given hierarchical level, taxa sharing similar sequences may not be resolved to lower taxonomic levels and thus goes to a higher taxonomic level, reducing resolution. This could be why a high proportion of reads (47.3%) mapped to GenBank were not useful for identification. In contrast, when using a national DNA reference database where only sequences of species found in the region are represented, such as the case for DNAmark, sequences are easily resolved to lower taxonomic levels with reduced or no misidentifications. Hence, national DNA reference databases provide better taxonomic resolution and accuracy. Depending on research needs, this necessitates the need for utilising a national curated DNA reference database (if available) rather than a global one in molecular studies, where accurate taxon identifications are crucial to meet research goals. However, if global DNA reference databases utilised can be of the same standards as national ones, matching reads to a global DNA reference database would be preferred as it would have allowed for the detection of contamination and species not expected in datasets [54]. For example, species that were found in the past, but are now locally extinct. The generated reference data from this study has other applications other than taxonomic assignment, such as for example the designing of primers or capture probes [45,46], understanding adaptive changes through evolutionary studies of plastid genome functions [89,90], and improving plant phylogeny resolution [91,92]. Of particular interest is the ability to design better primers and select taxon-specific barcodes, where genome skims of only a few taxa would be sufficient to provide enough data to improve upon the discriminatory power of current primers and barcodes used [13,34].

## CONCLUSIONS

We demonstrated that genome skimming of plants for the generation of reference data had a high rate of assembly success. Included in the generated reference data are some of the most important plant species for nature conservation and research in Denmark, and this study sets a good foundation for adding assemblies of other plant species in the near future. With 184 plant species sequenced, covering 45% of all Danish plant families, we have already added 101 cpGenomes and 6 nrDNA sequences of species that were previously not represented in global reference databases like GenBank. We further showcase through *in silico* metagenomics analysis of our generated reference data, the use of our national DNA reference data for molecular identification has better resolution and accuracy, resulting in more taxonomically informative reads as compared to GenBank. As this is the first national DNA reference database built for Danish species (DNAmark), it provides an important resource for researchers, particularly in molecular biodiversity monitoring studies where accuracy is key. Our study is a step forward in establishing a curated flora DNA reference database for plant molecular studies not just in Denmark, but on a global scale when combined with the reference data generated from other genome skimming studies [13,14,40]. These reference data can also be used in various other applications including designing better plant primers, contributing to the wider scientific community by providing blueprints for optimal utilisation of the genome skimming technique.

## SUPPLEMENTARY INFORMATION

### Supplementary Material 1

**Table S1:** Information on specimens used to build the DNAmark reference database

**Table S2:** Plant species used for sequencing depth exploration

**Table S3:** List of plants (assembled in the DNAmark project) used in each of the three in silico simulated datasets

**Table S4:** List of plants used in each in silico simulated dataset

**Table S5:** Comparison of database and species coverage used in the in silico simulation. Green: corresponding barcode sequence of species present in database; red: corresponding marker sequence of species absent in database

**Table S6:** Number of species with sequences not represented in GenBank contributed by the DNAmark project

**Table S7:** Species with sequences not represented in GenBank contributed by the DNAmark project

**Table S8:** Comparison of genome and barcode coverage between GenBank and DNAmark

**Table S9:** Information on genes recovered for each sample. Green: gene recovered; red: gene not recovered

**Table S10:** Output of sequencing depth exploration. Red: unassembled, green: assembled

**Table S11:** In silico application raw outputs – Taxonomic identification according to database used for each barcode and the number of reads mapped to the corresponding barcode

**Table S12:** In silico application outputs after applying identification parameters and barcode combination – Taxonomic identification according to the database used for each barcode and the number of reads mapped to the corresponding barcode. Red: Incorrect identification

**Table S13:** Recovery of plants used in each of the three in silico simulated datasets and the reference database used for taxonomic assignment (Red: not identified. green: identified)

**Table S14:** Taxonomic resolution for matches to GenBank or DNAmark reference database

### Supplementary Material 2

**Fig S1:** Proportion of sequenced samples with assembled chloroplast genomes (cpGenomes) and nuclear ribosomal DNA (nrDNA) sequences across the 54 plant families. N is the number of voucher specimens sequenced for each family.

**Fig S2:** Proportion of assembled a) cpGenomes and b) nrDNA sequences containing the five common barcodes (*rbcL*, *matK*, *trnL*, ITS1 & ITS2) used in taxonomic identification of plants.

## DECLARATIONS

### Availability of supporting data

Raw sequencing data are deposited in SRA under project number PRJNA607895. The assembled chloroplast genomes and nuclear ribosomal sequences have been uploaded to GenBank. The accession numbers (both GenBank and SRA) for each specimen can be found in Table S1_Supplementary Material 1. Files and scripts used for generating data will be made publicly available on the GigaScience Repository, GigaDB, upon acceptance of this manuscript.

## Abbreviations

*atpB*: adenosine triphosphate synthase subunit beta
BEMT: Blunt End Multi Tube
BOLD: Barcode of Life Data Systems
BP: base pairs
BSA: Bovine Serum Albumin
BWA: Burrows-Wheeler Alignment
CDS: Coding sequences
*COI*: cytochrome-c oxidase subunit 1
cpGenome: Chloroplast genome
DNAmark: Danish national DNA reference database
eDNA: environmental DNA
GATK: GenomeAnalysisTK
GBIF: Global Biodiversity Information Facility
HTS: high-throughput sequencing
ITS: internal transcribed spacers
LCA: lowest common ancestor
LSU rRNA: large subunit ribosomal ribonucleic acid
*matK*: maturase K
nrDNA: nuclear ribosomal sequences
ORG.Annot: Organelle Annotator
ORG.asm: ORGanelle ASseMbler
PE: paired-end
PVP: Polyvinylpyrrolidone
qPCR: quantitative PCR
*rbcL*: ribulose-bisphosphate carboxylase large chain
SD: standard deviation
sedaDNA: Ancient sedimentary DNA

## Consent for publication

Not applicable

## Competing interests

The authors have no competing interests.

## Funding

This research is funded by Aage V. Jensen Naturfond, to establish the Danish National DNA reference database (DNAmark). Additionally, PYSC is supported by the European Union Horizon 2020 research and innovation programme under grant agreement No 765000, H2020 MSCA-ITN-ETN Plant.ID network.

## Author contributions

PYSC, KB designed and implemented the research; CN coordinated sample collection on the broader DNAmark project; HHB, IH collected and identified samples; MER, SRR did the laboratory analysis; PYSC, AM planned and curated genome assembly pipelines; FL, EMRL did the assemblies; FL carried out sequencing depth exploration; PYSC conducted the *in silico* simulation; FL, EMRL did the statistical analysis. PYSC, EMRL, FL made the figures. PYSC led the writing of the manuscript with inputs from all authors. All authors contributed to the manuscript and approved the final version.

## Acknowledgement

We are grateful to the Natural History Museum of Denmark for the access to museum collections, assisting with specimen collections, and curating newly collected voucher specimens. Specifically, we thank Ole Seberg, Gitte Petersen, Nina Rønsted, Olof Ryding as well as Christian Lange. Furthermore, we wish to thank Ole Byrgesen, Louise Isager Ahl, Natalie Eva Iwanycki and Sam Bruun-Lund for their help and support with the collection of specimens. We would also like to thank the DNAmark committee for their invaluable inputs, specimen collection, and taxonomic identification of voucher specimens. We further thank the Danish National High-throughput DNA Sequencing Center for generating the Illumina data.

## Notes

### Competing Interest Statement

The authors have declared no competing interest.

